# Minimal Gene Signatures Enable High-Accuracy Prediction of Antibiotic Resistance in *Pseudomonas aeruginosa*

**DOI:** 10.1101/2025.04.29.651273

**Authors:** Nabia Shahreen, Syed Ahsan Shahid, Mahfuze Subhani, Adil Al-Siyabi, Rajib Saha

## Abstract

Antimicrobial resistance (AMR) in Pseudomonas aeruginosa poses a critical global health challenge, with current diagnostics relying on slow, culture-based methods. Here, we present a ML framework leveraging transcriptomic data to predict antibiotic resistance with high accuracy. We applied a genetic algorithm to 414 clinical isolates to identify minimal, highly predictive gene sets (∼35–40 genes) distinguishing resistant from susceptible strains for meropenem, ciprofloxacin, tobramycin, and ceftazidime. Automated ML classifiers trained on these sets achieved accuracies of 96–99% on test data (F1 scores: 0.93–0.99), surpassing clinical deployment thresholds. Multiple distinct, non-overlapping gene subsets exhibited comparable performance, indicating that resistance acquisition broadly impacts the expression of diverse regulatory and metabolic genes. Comparison with known resistance markers from CARD and operon annotations revealed a substantial number of previously unannotated clusters, highlighting significant knowledge gaps in current AMR understanding. Mapping these genes onto independently modulated gene sets (iModulons) revealed transcriptional adaptations across diverse genetic regions. Overall, this study presents a streamlined machine-learning workflow for transcriptomic data and offers a pathway toward rapid diagnostics and personalized treatment strategies against AMR.

## Introduction

Antimicrobial resistance has emerged as one of the most urgent threats to global public health, undermining the effectiveness of existing treatments and increasing the risk of untreatable infections. The World Health Organization (WHO) has identified AMR as one of the top ten global health threats, projecting that, without intervention, it could lead to 10 million deaths annually by 2050 (World Health Organization, 2019). Among the most concerning pathogens is *P. aeruginosa*, a gram-negative opportunistic bacterium responsible for severe infections such as pneumonia, urinary tract infections, and bacteremia, particularly in immunocompromised patients (Horcajada *et al*, 2019; Qin *et al*, 2022; Pang *et al*, 2019). The threat posed by this bacterium is exacerbated by its intrinsic resistance mechanisms, including efflux pumps and reduced outer membrane permeability, as well as its ability to rapidly acquire new resistance determinants, leading to the emergence of multidrug-resistant (MDR) and even pan-resistant strains (Breidenstein *et al*, 2011; Lister *et al*, 2009).

Despite the escalating threat, clinical practice still relies primarily on culture-based antibiotic susceptibility testing, which, while reliable, can require 48–72 hours to yield results (van Belkum *et al*, 2019). This delay necessitates empirical treatment with broad-spectrum antibiotics, which may be ineffective and can further drive resistance (Tacconelli *et al*, 2018). Moreover, culture-based assays provide limited insight into the genetic or molecular drivers of resistance, offering little guidance for targeted interventions (Dyar *et al*, 2017).

In recent years, high-throughput sequencing, particularly transcriptomic profiling, has enabled a more fine-grained view of resistance by capturing the gene expression that underpin survival under antibiotic pressure (Lázár *et al*, 2018; Alsiyabi *et al*, 2024; Khaledi *et al*, 2016; Suzuki *et al*, 2014). Transcriptomic data provides a snapshot of the cellular state under antibiotic pressure, revealing pathways and regulatory networks that contribute to survival. This approach holds promise for identifying biomarkers of resistance, enabling earlier and more precise diagnostics (Bhattacharya *et al*, 2017; Wang *et al*, 2022; Khaledi *et al*, 2016). However, leveraging transcriptomic data for AMR surveillance and prediction remains challenging due to the high dimensionality of the data, which complicates the identification of relevant features for accurate and interpretable predictions (Topçuoğlu *et al*, 2020).

One promising avenue for overcoming these challenges is machine learning, which can handle large omics datasets and uncover complex patterns relevant to resistance phenotypes (Libbrecht & Noble, 2015). Prior studies have demonstrated the feasibility of ML-driven AMR prediction using genomic and transcriptomic data, highlighting the potential for faster and more precise diagnostics (Macesic *et al*, 2020). However, a major challenge remains in identifying minimal yet robust gene subsets that preserve predictive accuracy while improving interpretability and reducing computational costs. In a notable study, Khaledi et al, 2020 demonstrated that integrating single nucleotide polymorphisms (SNPs), gene presence/absence (GPA), and transcriptomic expression data enabled predictive modeling of antibiotic resistance with sensitivity and predictive values between 0.81 and 0.95 across four antibiotics. However, their reliance on a high-dimensional feature space (up to 93 markers per antibiotic) and mixed-data approaches limit clinical scalability due to cost, interpretability, and generalizability constraints. Additionally, their pipeline required manual feature selection and hyperparameter tuning, lacking a fully optimized and automated feature selection mechanism.

To address these limitations, we implemented a hybrid methodology combining genetic algorithm (GA)-based feature selection with automated ML (AutoML), leveraging transcriptomic data from 414 *P. aeruginosa* clinical isolates (Figure 1, Appendix Figure S1). By iteratively evolving gene subsets, GA identified minimal, highly informative features, which were then used to train ML models with accuracies of 0.96–0.99 on held-out test data. Despite minimal overlap among gene subsets, each yielded similarly high predictive power, suggesting a pervasive, multifactorial transcriptomic signature of resistance. To resolve whether these subsets shared underlying biological processes beyond direct gene overlap, we employed three complementary analyses. First, we compared GA-selected genes to known resistance determinants in the CARD, examining the extent to which recognized resistance markers drove the predictions (Alcock *et al*, 2023). Second, we explored operon-level organization to evaluate whether co-transcribed gene clusters were consistently selected, revealing potential regulatory “hotspots” (Winsor *et al*, 2016; Cao *et al*, 2019). Finally, we mapped the subsets to iModulons, co-regulated gene modules derived from Independent Component Analysis (ICA), to elucidate higher-order transcriptional control mechanisms associated with resistance (Lee, 1998; Sastry *et al*, 2021; Rajput *et al*, 2022). Our results show that gene expression patterns predictive of AMR in *P. aeruginosa* extend beyond canonical resistance genes, suggesting that diverse transcriptional responses may characterize the resistant phenotype. While some GA-selected loci mapped to well-characterized efflux pumps or β-lactamase operons, many fell outside conventional AMR annotations, pointing to underexplored determinants. Operon-level analysis identified recurrent co-transcribed clusters involved in osmotic stress, iron acquisition, and various metabolic pathways, while iModulon mapping revealed a convergence on transcriptional programs governing oxidative stress responses, DNA repair, efflux regulation, and ribosomal function. Collectively, these data highlight that AMR phenotypes correlate with transcriptomic patterns spanning diverse genetic loci, including both isolated resistance genes and genes implicated in broader cellular processes.

**Figure 1:**
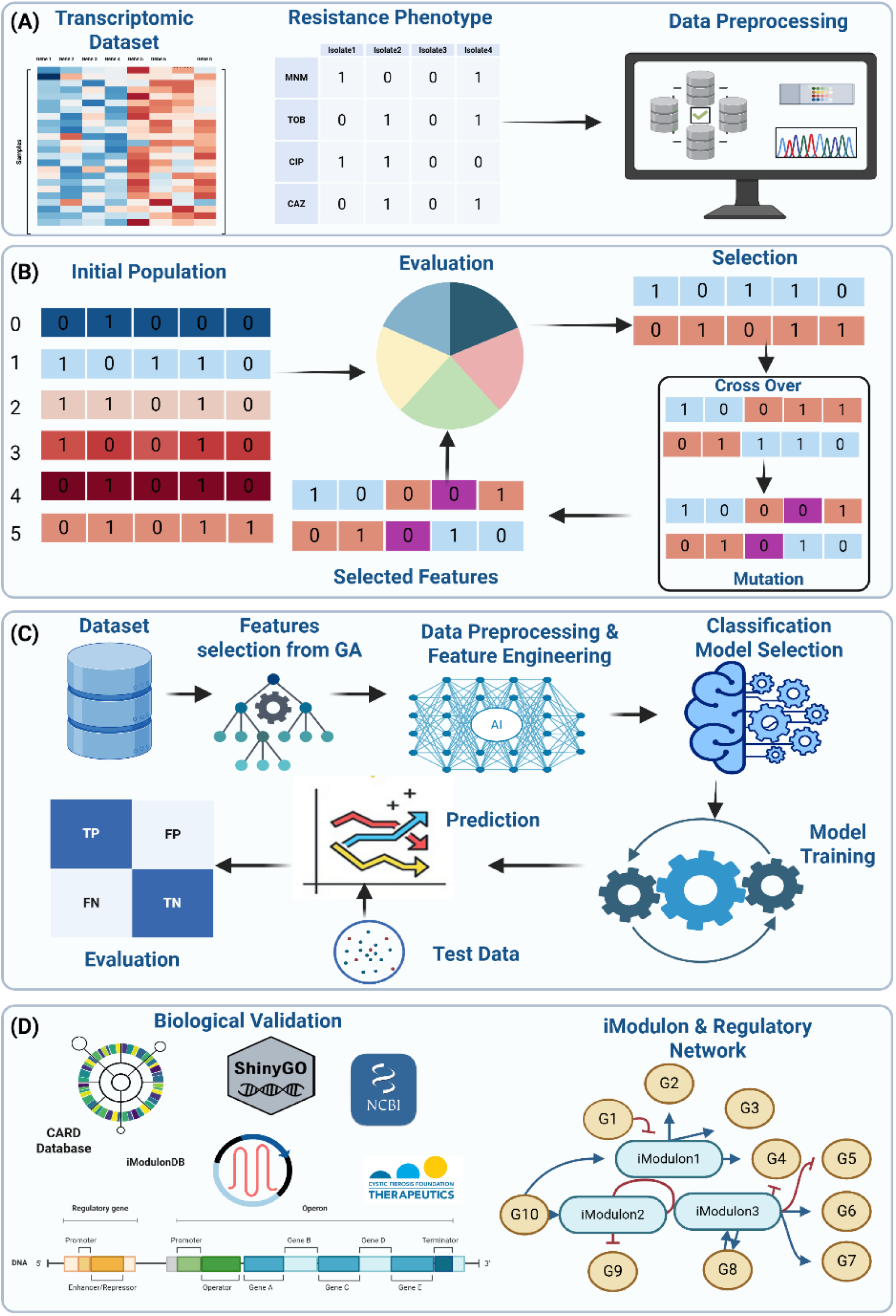
Overall pipeline for the GA–AutoML workflow and biological validation in *P. aeruginosa*. (A) Incorporation of RNA-seq data from 414 clinical isolates alongside antibiotic phenotypes and initial data processing. (B) Application of a genetic algorithm that iteratively identifies minimal, high-accuracy gene subsets through population initialization, selection, crossover, and mutation steps. (C) Integration of these GA-selected features into data preprocessing, classification, model selection, and final performance evaluation under an automated machine-learning pipeline. (D) Post hoc validation of selected genes through multiple external resources (CARD, Pseudomonas Genome DB, ShinyGO, NCBI, iModulonDB), focusing on their operon assignments, potential regulatory networks, and broader functional context (Alcock *et al*, 2023; Winsor *et al*, 2016; Ge *et al*, 2020; O’Leary *et al*, 2024; Sastry *et al*, 2024).

Overall, in this study, we present a multi-scale framework that integrates transcriptomic profiling, evolutionary feature selection, and in-depth biological network analysis to enhance AMR diagnostics. By pinpointing compact yet functionally enriched gene sets linked to specific operons and regulatory modules, our approach offers a balance of predictive accuracy, interpretability, and clinical feasibility. These findings underscore the value of system-level perspectives on AMR and support the development of next-generation diagnostics and personalized therapeutic strategies.

## Results

### Automated ML and Genetic Algorithm Uncover Minimal Yet High-Performing Gene Sets

Predicting antibiotic resistance from gene expression in *P. aeruginosa* requires a balance between accuracy and interpretability, as high-dimensional transcriptomic data presents computational and clinical challenges. To address these constraints, we developed a hybrid GA-AutoML pipeline designed to systematically identify minimal, highly predictive gene subsets while optimizing classification performance. Initially, AutoML alone, using all available 6,026 genes, yielded strong baseline models with accuracy up to 0.9 and F1-scores up to 0.88 on a holdout set for each antibiotic: meropenem (MNM), ciprofloxacin (CIP), tobramycin (TOB), and ceftazidime (CAZ). Although these results demonstrate high predictive accuracy, reliance on the entire transcriptome poses substantial computational and sequencing challenges, limiting routine clinical adoption.

To address the challenge of high dimensionality in transcriptomic data, we employed GA (Mirjalili, 2019) to systematically identify compact gene subsets capable of accurately predicting antibiotic resistance phenotypes. The process began with a randomly initialized population of 40-gene subsets and iteratively refined them over 300 generations per run. In each generation, candidate subsets were evaluated using support vector machines (SVM) and logistic regression (LR), with classification performance assessed through ROC-AUC and F1-score metrics (see Methods). High-performing subsets were preferentially retained and recombined using selection, crossover, and mutation operations, ensuring continued exploration of diverse gene combinations. This process was repeated independently for 1,000 runs per antibiotic, resulting in a broad array of high-performing but largely non-overlapping feature sets.

Rather than converging on a single fixed subset, the GA produced thousands of distinct gene combinations, each achieving strong predictive performance. We observed that certain genes were consistently selected across many independent runs, suggesting their robust association with resistance phenotypes. To construct clinically practical and interpretable models, we generated consensus gene sets by ranking all genes based on their frequency of selection across GA iterations. These top-ranked genes, representing the most repeatedly selected and biologically plausible features, were then used to train final classifiers for each antibiotic.

The average performance of these consensus-based models was comparable to or exceeded full-transcriptome classifiers, with test set accuracies of ∼99% for MNM and CIP, and ∼96% for TOB and CAZ (Figure 2A, Dataset EV1). These compact classifiers, typically comprising 35–40 genes per antibiotic, offer an interpretable and computationally efficient alternative to transcriptome-wide approaches.

**Figure 2.**
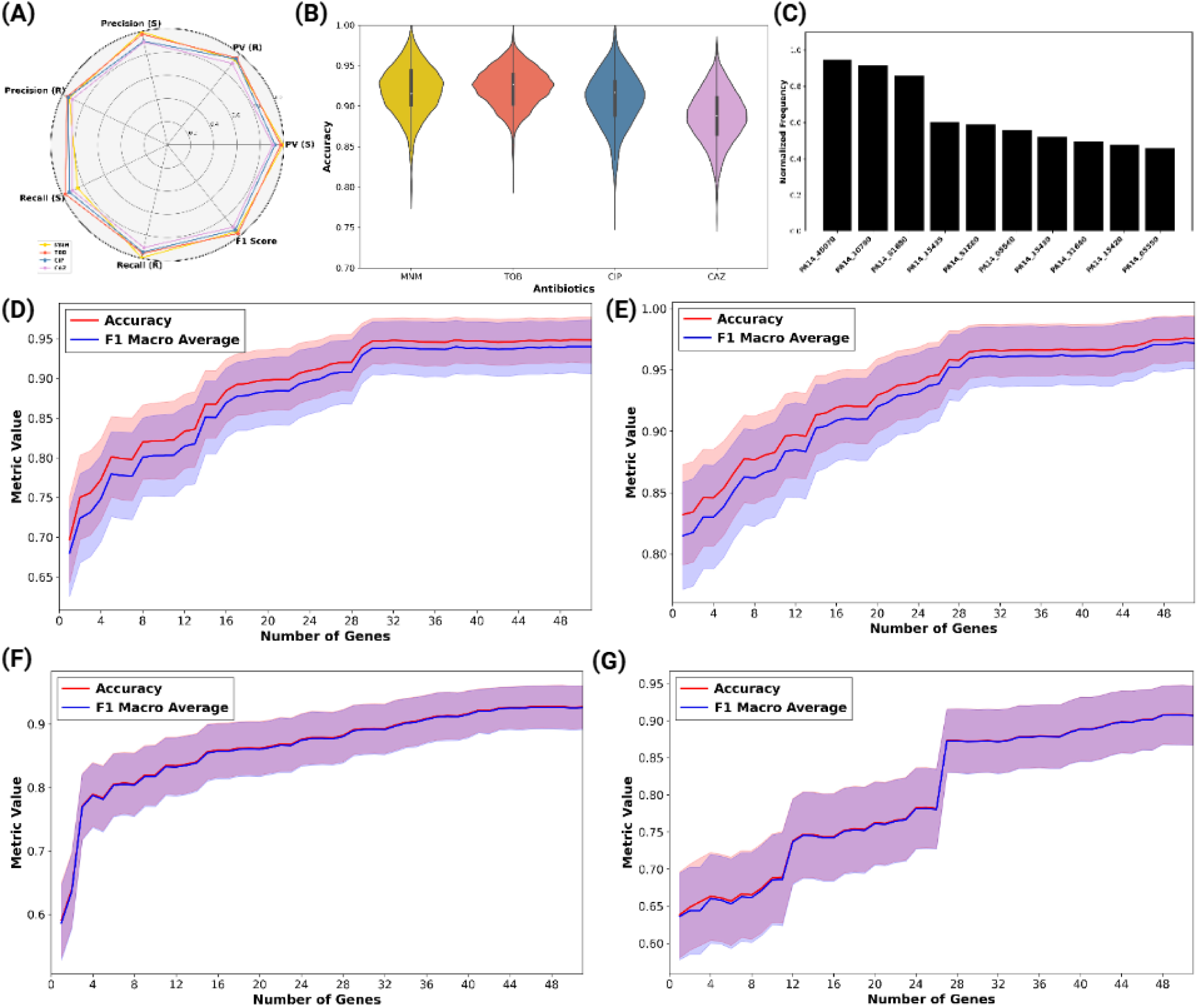
Classification performance and feature set optimization across four antibiotics. (A) Radar chart displaying precision, recall, predictive value (PV), and F1-score (0–1 range) for susceptible (S) and resistant (R) classes in the top-performing feature sets for each antibiotic. (B) Violin plots illustrating accuracy distribution across 1,000 genetic algorithm iterations, with median accuracy (dotted line), interquartile range (shaded area), and density. MNM and CAZ exhibit slightly broader variability, while TOB and CIP demonstrate narrower distributions around high accuracies. (C) Representation of the top 10 genes that were most repeated in GA runs across all antibiotics. (D-G) Classifier performance (accuracy and F1 macro-average) as a function of the number of most frequently selected genes used for model training, shown separately for MNM (D), TOB (E), CIP (F), and CAZ (G). Lines represent mean performance across cross-validation runs; shaded areas denote standard deviation. Performance improvements plateau at ∼35–40 genes, supporting the use of compact consensus feature sets for accurate and stable prediction of resistance phenotypes.

To evaluate model robustness, we examined the accuracy distributions across 1,000 GA iterations. CIP and TOB classifiers showed narrower variability and higher median performance, while MNM and CAZ demonstrated broader, yet still high-performing, distributions (see Figure 2B). We also ranked genes by their selection frequency and plotted the top 10 genes that were most consistently selected across antibiotics (Figure 2C).

We further assessed how model performance varied with the number of top-ranked genes used for classification. For each antibiotic, performance plateaued after inclusion of ∼35–40 genes, justifying the selection of the number of features (Figure 2D–G). This plateau was observed for both accuracy and macro-average F1-score, underscoring that a limited number of frequently selected genes suffices to capture the transcriptomic signature of resistance.

To further enhance interpretability, we filtered iteration-specific subsets to retain ≥80% annotated genes. This ensured that the selected signatures remained both biologically informative and clinically actionable. The GA framework thus provided two complementary outputs: (1) iteration-specific subsets for assessing variability and classification performance, and (2) consensus-ranked gene lists for biological interpretation and downstream regulatory analysis (see Methods).

Collectively, our integrated GA–AutoML approach identified minimal yet highly predictive transcriptomic signatures across multiple antibiotics. The repeated selection of previously unannotated genes highlights underexplored regulatory or metabolic features of resistance, offering avenues for future biological investigation.

### GA-Derived Subsets Reveal Limited CARD Overlap and Highlight Uncharacterized Genes

To determine whether our GA-AutoML-derived genes correspond to known antibiotic resistance markers or represent previously unrecognized determinants, we compared the top-performing antibiotic-specific gene subsets to genes annotated in the Comprehensive Antibiotic Resistance Database (CARD) (Alcock *et al*, 2023). Across all antibiotics, this analysis revealed that only a minor fraction (2–10%) of our predictive gene subsets overlapped with established AMR genes listed in CARD (Dataset EV2), indicating substantial novelty within our gene signatures (Figure 3).

**Figure 3:**
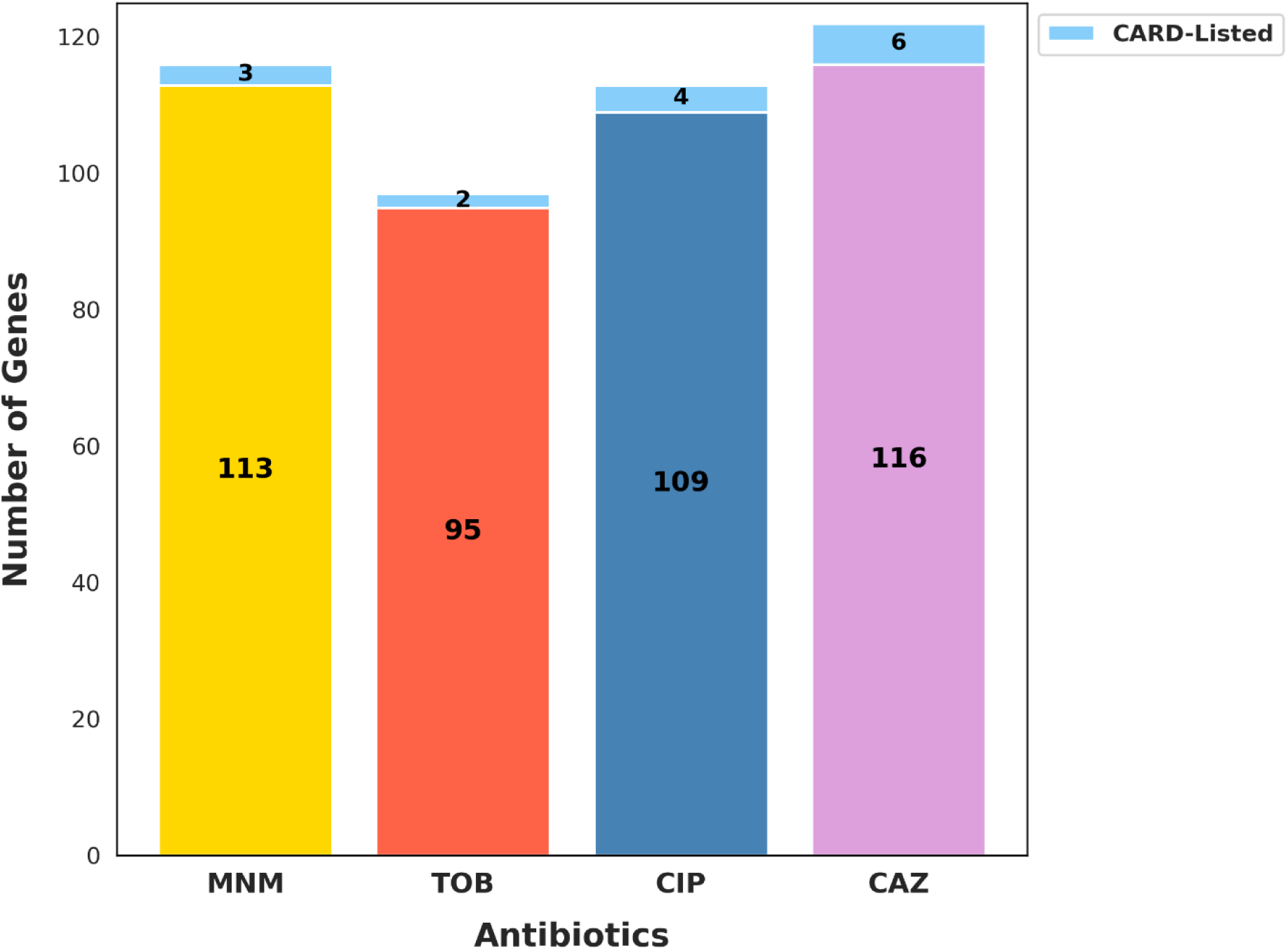
GA-Derived Subsets Reveal Minimal Overlap and Predominantly Novel AMR Genes Relative to CARD. Stacked bar plot comparing the number of GA-selected genes overlapping with the CARD Database versus those absent from CARD across the four antibiotics: MNM, TOB, CIP, and CAZ. For each antibiotic, the total number of unique genes identified from high-performing GA-derived subsets is partitioned into CARD-listed and novel categories. The small size of the known segment in most cases underscores the substantial novelty captured by our approach.

For MNM, the overlap with CARD-annotated genes was approximately 3–5%, involving efflux pump-related loci *mexA* and *mexB*, both selected in >50% of GA iterations. These genes are well-documented contributors to resistance via antibiotic efflux (Lorusso *et al*, 2022; Poole, 2005). Interestingly, the most consistently selected gene by GA, *gbuA*, chosen in approximately 95% of iterations, is absent from CARD. *GbuA* encodes guanidinobutyrase, an enzyme in arginine catabolism (Nakada & Itoh, 2002). Upregulation of *gbuA* in meropenem-resistant isolates may serve as a compensatory biomarker for impaired arginine uptake linked to loss of function mutations in *OprD*, a porin gene critical for carbapenem entry (Li *et al*, 2012). While *gbuA* itself is not directly implicated in resistance mechanisms, its strong association with resistant phenotypes highlights its utility as a diagnostic marker.

In predicting TOB resistance, only a limited overlap (2–5%) was found between GA-derived genes and CARD annotations. The gene PA14_15435, encoding a cyclic di-GMP-specific phosphodiesterase, was prominently selected (∼80% iterations) but is not annotated in CARD. Its activity may influence biofilm formation, motility, and antibiotic penetration, indirectly affecting aminoglycoside susceptibility (Valentini & Filloux, 2016). Other highly selected genes (PA14_15430, PA14_15420, PA14_15410) correspond to transposon-associated elements linked to a genomic island carrying mercury resistance genes (Partridge *et al*, 2018). Although these do not directly confer resistance, their repeated selection indicates that genomic context or co-selection events might mark isolates adapted to aminoglycoside stress. Known aminoglycoside resistance genes, such as *amrB* (PA14_60860), were less frequently selected (∼30%), suggesting that despite their direct resistance roles, they exhibit weaker transcriptomic signatures distinguishing resistant strains.

For CIP, the most frequently selected genes by GA were predominantly absent from CARD annotations as well. PA14_61650 (*pagL*, >70% iterations), encoding lipid A 3-O-deacylase, modifies LPS structure, potentially influencing antibiotic permeability (Rutten *et al*, 2006). PA14_31640 (glyoxalase I homolog, ∼40% iterations) may mitigate oxidative stress induced by ciprofloxacin treatment. PA14_36990 encodes another cyclic di-GMP phosphodiesterase, influencing biofilm dynamics and possibly efflux mechanisms indirectly affecting fluoroquinolone susceptibility (Valentini & Filloux, 2016). Meanwhile, genes known for quinolone resistance (e.g., *mexA, mexB, gyrA*), identified in CARD, were infrequently selected (10–20%), likely due to subtler transcriptomic differences.

In CAZ resistance prediction, significant overlap with CARD was observed due to the frequent selection (>90%) of ampC (PA14_10790), a known β-lactamase (Livermore, 1995). However, PA14_33680 (*fpvA*, >40% iterations), encoding the ferripyoverdine receptor, was repeatedly selected despite not being in CARD. Elevated *fpvA* expression has been correlated with ceftazidime resistance, possibly linking iron acquisition responses with antibiotic resistance phenotypes (Ramsay *et al*, 2023).

Overall, our analysis underscores that GA-selected genes frequently fall outside conventional AMR annotations, likely because they show pronounced transcriptomic differences between resistant and susceptible isolates, rather than directly conferring resistance. Many of these genes play indirect roles, such as metabolic compensation, membrane modification, stress responses, or biofilm regulation. These results suggest that many resistance-associated transcriptomic signatures fall outside current AMR annotations, reflecting that conventional databases may not capture the full landscape of expression-based adaptations observed in resistant strains. While not direct resistance determinants, such genes may serve as robust biomarkers or proxies for resistant phenotypes, meriting further experimental characterization.

### Operon-Level Analysis Reveals Limited Functional Overlap in Predictive Gene Sets

In addition to genes lacking prior association with resistance mechanisms, our GA-AutoML pipeline generated multiple gene subsets per antibiotic, each containing approximately 35-40 genes, as predictive models. These subsets achieved high predictive accuracy (∼96–99%) despite minimal gene overlap (∼5–8%) within each antibiotic category (Figure 4 A-D). This unexpected observation prompted us to examine whether these subsets share common biological functions at the operon level. An operon is a group of genes in prokaryotes (bacteria) that are transcribed together as a single mRNA molecule (Salgado *et al*, 2000). By analyzing which operons the selected genes belonged to, we asked whether different subsets might still be capturing similar biological responses, even if the specific genes varied.

**Figure 4:**
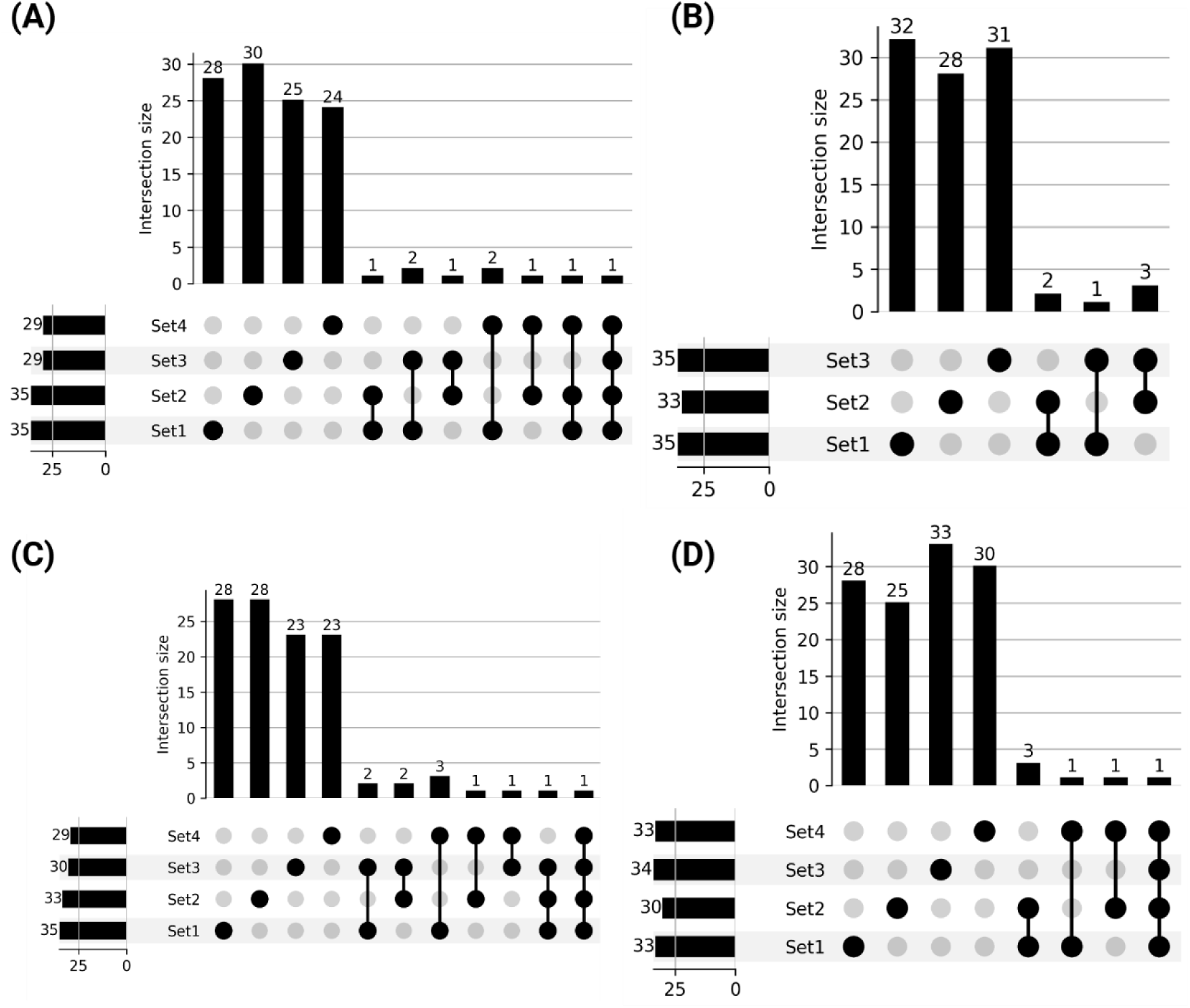
Minimal Overlap Among Multiple High-Accuracy GA-Selected Gene Subsets. UpSet plots illustrating the pairwise and higher-order intersections among the top GA-selected gene subsets for MNM (A), TOB (B), CIP (C), and CAZ (D), respectively. Each UpSet plot underscores the limited overlap across these equally high-accuracy subsets, highlighting distinct routes to resistance.

Several antibiotic-specific operons were indeed consistently represented across multiple predictive gene subsets (Dataset EV3). For MNM, two operons consistently stood out: the well-characterized mexAB-oprM operon, encoding a multidrug efflux pump system directly involved in antibiotic efflux (Poole, 2005), and the gbuA operon, encoding guanidinobutyrase and related enzymes (Figure 5A). The frequent identification of mexAB-oprM aligns with its established role in direct antibiotic resistance, particularly through reducing intracellular meropenem concentrations (Avakh *et al*, 2023). The recurrent selection of the gbuA operon suggests its involvement in broader metabolic adaptations, potentially compensating for nutrient uptake disruptions such as those resulting from impaired porin function (e.g., OprD deficiency), which is a known determinant of carbapenem resistance (Wang *et al*, 2025; Jagmann *et al*, 2016).

**Figure 5:**
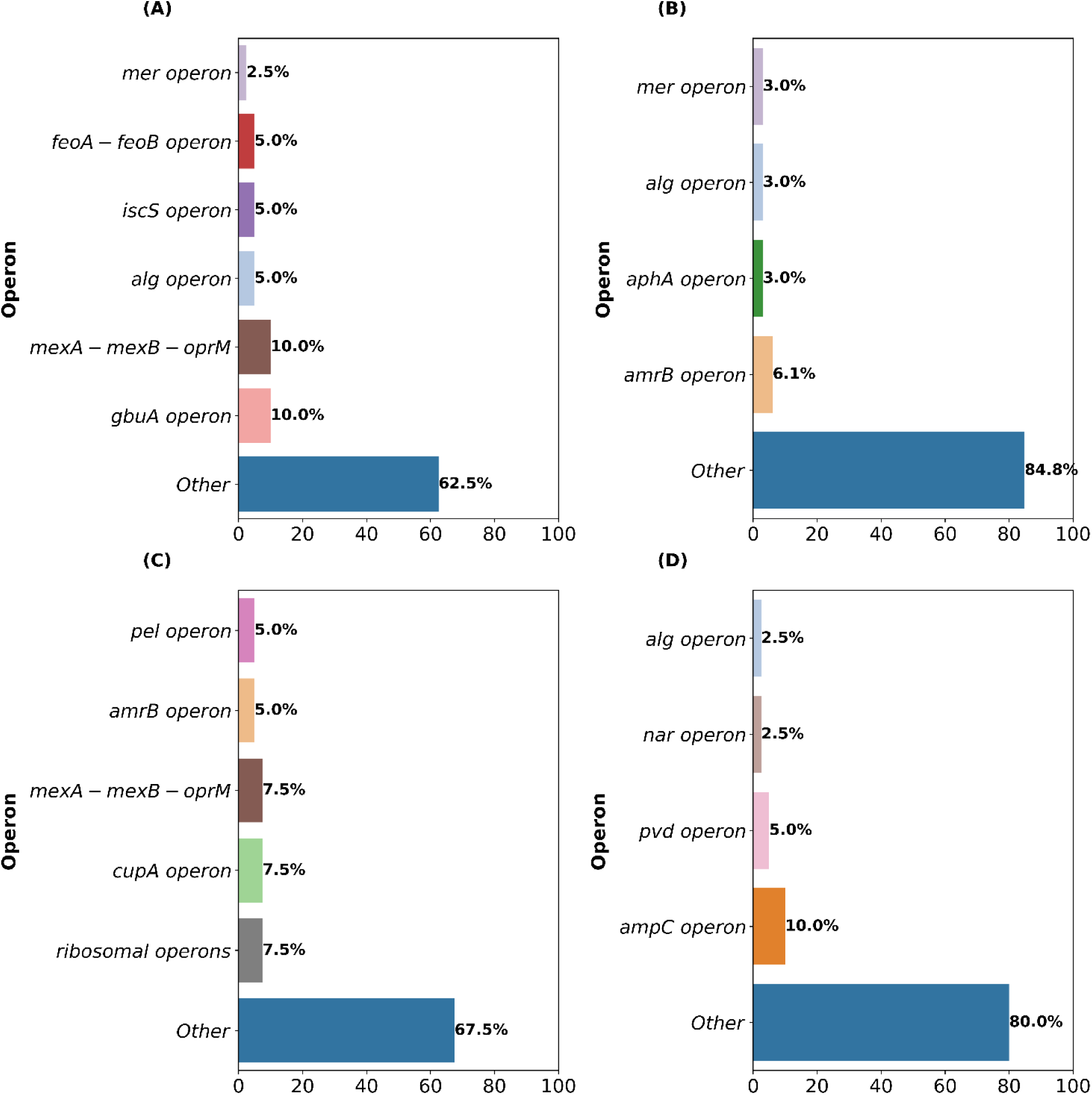
Recurrent and Dispersed Operon Involvement Across GA-Predictive Gene Sets. Bar plots show how often genes from specific operons were selected across multiple high-performing subsets for each antibiotic: MNM (A), TOB (B), CIP (C), and CAZ (D). While known resistance operons such as *mexAB-oprM* and *ampC* were repeatedly selected, most predictive genes came from diverse operons, highlighting that resistance-associated transcriptomic signatures involve diverse genetic loci beyond classical resistance operons.

For TOB, operon-level recurrence was more limited, highlighting diverse, isolate-specific strategies potentially reflecting antibiotic exposure history or genomic context. The aminoglycoside efflux operon amrAB was identified in two subsets, consistent with its direct role in antibiotic expulsion. Interestingly, the mercury resistance operon (mer) and alginate biosynthesis (alg) operon each appeared in single subsets (Figure 5B). Although mercury resistance (mer operon, e.g., gene PA14_15435) is not directly linked to antibiotic resistance, its selection could indicate a broader cellular stress response or biofilm adaptation influencing aminoglycoside penetration or susceptibility. Similarly, the alg operon’s selection may reflect the known association between biofilm formation and reduced antibiotic susceptibility (Uruén *et al*, 2020).

In CIP resistance prediction, the efflux operon mexAB-oprM appeared frequently, underscoring its well-documented role in fluoroquinolone efflux (Piddock, 2006) (Figure 5C). Additionally, the pel operon (biofilm polysaccharide production), cupA operon (fimbrial adhesion), and ribosomal operons emerged recurrently, suggesting additional layers of antibiotic response including biofilm formation and stress-induced translational adaptation, potentially mitigating ciprofloxacin-induced DNA damage (Friedman & Kolter, 2004). The diverse representation of operons involved in biofilm formation (pel) and adhesion (cupA) highlights broader physiological adaptations affecting antibiotic permeability and tolerance (Trastoy *et al*, 2018; Meissner *et al*, 2007).

Meanwhile, CAZ exhibited a distinct operon signature heavily dominated by the β-lactamase-encoding ampC-ampR operon (Glen & Lamont, 2021), selected across all subsets (Figure 5D). This was consistent with its direct enzymatic role in β-lactam hydrolysis. Secondary selection included iron scavenging (pvd operon) and anaerobic respiration operons (nar operon) (Punchi Hewage *et al*, 2020), reflecting metabolic and nutritional adaptations potentially associated with resistance phenotypes, particularly under nutrient-limited infection conditions where antibiotic tolerance may increase.

Critically, however, despite the consistent selection of certain operons, most predictive genes (approximately 70–85% across antibiotics) did not map to these recurrent operons. Instead, these genes were broadly dispersed throughout the genome, spanning various unrelated functional groups and metabolic pathways. This extensive dispersion suggests that antibiotic resistance-associated transcriptomic signatures capture generalized transcriptional responses rather than being strictly limited to specific operon-based regulatory adaptations.

The broad genomic distribution of predictive genes further emphasizes that antibiotic resistance involves not only direct antibiotic neutralization (e.g., efflux pumps, antibiotic-modifying enzymes) but also indirect responses including metabolic shifts, stress responses, biofilm dynamics, and cell-envelope modifications. These diverse transcriptomic adaptations reflect the multi-layered complexity underlying antibiotic resistance in *P. aeruginosa*.

### iModulon Mapping Reveals Regulatory Convergence and Strain-Specific Adaptations

To further explore the regulatory basis of these resistance signatures, we examined whether these diverse operons may be orchestrated by shared regulatory modules, as identified through iModulon mapping. First, we mapped the feature sets to published iModulons of *P. aeruginosa* PAO1, which represent co-regulated gene groups identified through ICA (Rajput *et al*, 2022). Approximately 40–60% of feature set genes consistently mapped to known iModulons, forming a clear regulatory framework. Conversely, 30–50% remained unmatched, with 12% of these corresponding to PA14-specific genes absent in PAO1. These results underscore the regulatory complexity and strain-specific adaptations underpinning AMR mechanisms.

Mapping the feature sets to PAO1 iModulons revealed that distinct gene subsets converged on shared regulatory modules associated with critical cellular processes (Dataset EV4). Several feature sets mapped to iModulons involved in oxidative stress adaptation, a common bacterial response to antibiotic-induced damage. These iModulons include genes encoding catalases, superoxide dismutases, and other antioxidants, which help mitigate the effects of reactive oxygen species generated by antibiotics (Higazy *et al*, 2024). Similarly, multiple feature sets were enriched in iModulons associated with efflux pump systems, such as the MexAB-OprM and MexCD-OprJ operons. These systems are known to confer resistance by expelling antibiotics from the cell, reducing intracellular drug concentrations to sub-lethal levels (Lorusso *et al*, 2022).

The feature sets for TOB and other antibiotics frequently mapped to iModulons regulating ribosomal proteins and translation machinery, reflecting the role of protein synthesis inhibition in antibiotic action (Arenz & Wilson, 2016). For CIP, the feature sets consistently mapped to iModulons involved in DNA repair and recombination, aligning with the mechanism of action of fluoroquinolones, which target DNA gyrase and topoisomerase IV. These iModulons include genes encoding RecA and other DNA repair proteins, which are critical for surviving CIP-induced DNA damage.

To investigate strain-specific regulation, we performed a complementary ICA using 414 transcriptomes of clinical PA14 isolates, yielding a PA14-specific iModulon set (Dataset EV5). Mapping the GA-selected gene subsets onto this PA14 compendium showed high coverage for each antibiotic. For MNM, the four predictive subsets exhibited 83–88% alignment with PA14 iModulons (Fig. 6A), predominantly mapping to modules related to metabolism and nutrient adaptation, general stress response, efflux mechanisms, and cell envelope integrity. The CIP-associated subsets demonstrated 81–83% mapping (Fig. 6B), with prominent allocation to metabolic adaptation, DNA repair, and general stress response iModulons. In the CAZ subset, 80– 82% of genes mapped primarily to modules associated with metabolic pathways, cell envelope remodeling, and stress responses (Fig. 6C). Finally, the TOB subsets exhibited 79–81% mapping (Fig. 6D), most strongly associated with metabolic regulation, ribosomal functions, and stress adaptation. Across all antibiotics, the “Metabolism and Nutrient Adaptation” category consistently accounted for the largest fraction of mapped genes (Fig. 6A–D), reflecting the broad impact of antibiotic exposure on global metabolic rewiring beyond direct drug-target interactions.

**Figure 6:**
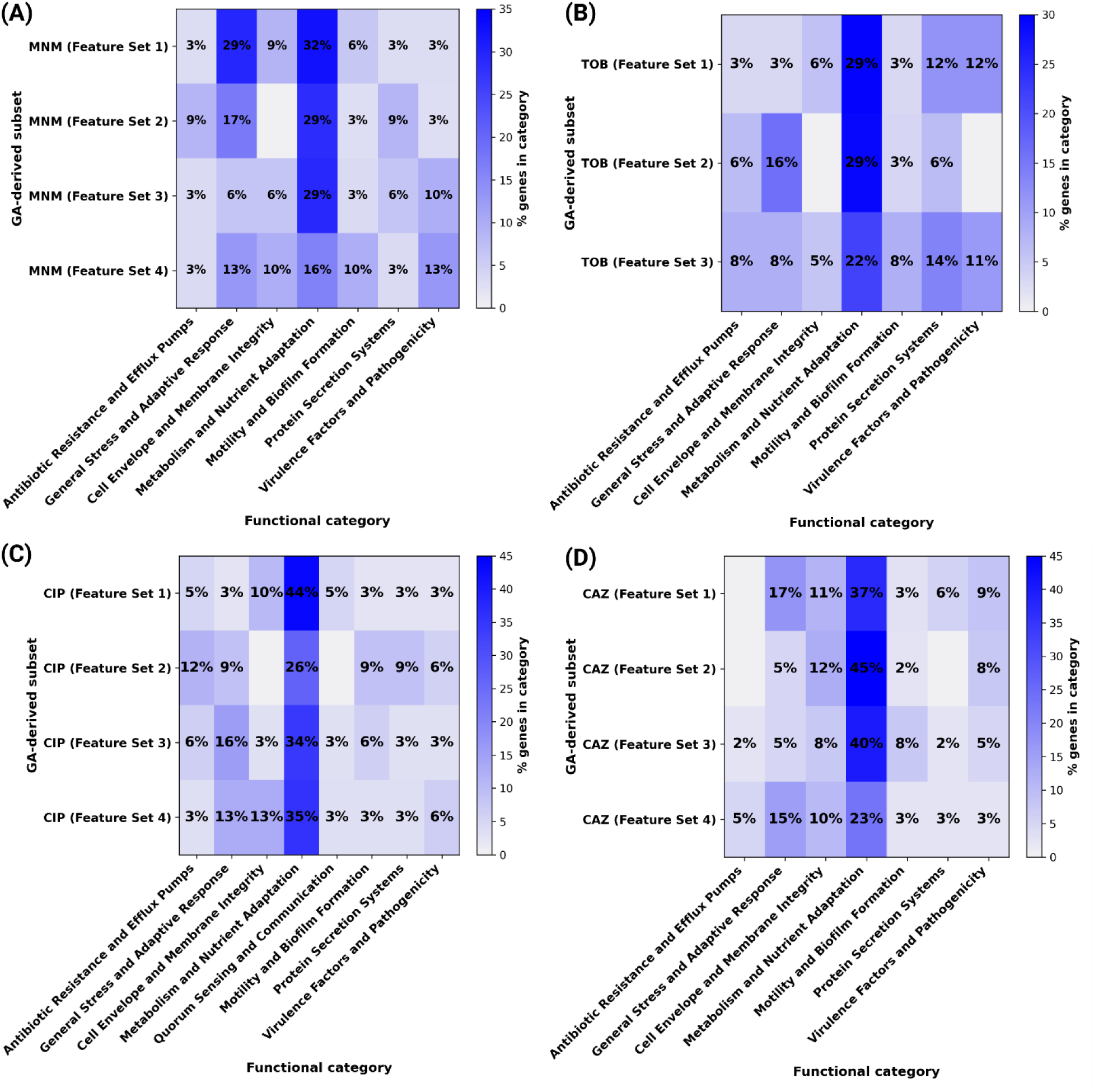
Functional allocation of GA-selected genes within PA14 iModulons. Heat-maps display the percentage of genes in each high-accuracy feature set (rows) that fall into specific PA14 iModulon functional classes (columns). for MNM (A), TOB (B), CIP (C), and CAZ (D). The color scale indicates the percentage of genes assigned to individual functional categories, showing that seemingly distinct gene sets converge on shared regulatory modules such as efflux, general stress, and metabolic adaptation.

These PA14-based findings mirror the regulatory themes observed with the PAO1 analysis, reinforcing the recurrent involvement of SOS-mediated repair, efflux systems, oxidative stress responses, ribosomal regulation, and envelope remodeling in antibiotic resistance across strains. Notably, several PA14-specific iModulons lacked direct counterparts in PAO1, including modules linked to heavy-metal detoxification and type VI secretion systems, highlighting strain-specific regulatory architectures that may contribute to multifactorial resistance strategies.

## Discussion

In this study, we demonstrated a successful automated approach for antibiotic susceptibility profiling in *P. aeruginosa* using only transcriptomic data. By applying a genetic algorithm to select informative features and an AutoML framework to train classifiers, we identified minimal gene expression signatures that achieved accuracies approaching 99% on a holdout set. Notably, this level of performance is comparable to existing genome-based approaches but relies on a markedly smaller set of transcriptomic features (Su *et al*, 2019; Khaledi *et al*, 2016, 2020; Suzuki *et al*, 2014). These results demonstrate the feasibility of building efficient, clinically actionable diagnostics based on compact transcriptomic signatures.

Many of the identified biomarkers include genes previously implicated in resistance, lending biological credibility to our approach. For example, *gyrA*, the primary target of fluoroquinolones (Akasaka *et al*, 2001), was recurrently selected in ciprofloxacin resistance signatures. Likewise, *ampC* and *oprD*, both of which are well-established β-lactam resistance determinants (Freed & Hanson, 2024), were frequently selected in ceftazidime and meropenem classifiers. Multidrug efflux pump components such as *mexA*, *mexB*, and *oprM* were also commonly identified across multiple antibiotics. The recurrence of these canonical resistance markers serves as internal validation and confirms that our feature selection strategy recovers biologically meaningful predictors.

Importantly, several highly ranked genes have not been previously annotated as resistance determinants. These include loci associated with stress adaptation, cell envelope remodeling, alginate biosynthesis, and metabolic regulation. Similar to earlier findings (Torrens *et al*, 2019; Kuper *et al*, 2024), our results suggest that resistant strains adopt broader physiological and regulatory adaptations beyond direct drug-target modifications. These genes may not themselves confer resistance but may act as transcriptional proxies or participate in compensatory mechanisms that enhance bacterial fitness under antibiotic stress. Their repeated selection across independent GA runs highlights their potential utility as biomarkers, even if they are not mechanistically causative.

Interestingly, the gene subsets identified for each antibiotic showed minimal overlap, indicating that multiple, equally predictive transcriptomic signatures can distinguish resistant from susceptible strains. While each subset included a small number of genes associated with known regulatory modules, such as oxidative stress response, SOS-mediated DNA repair, and efflux regulation, these represented only a fraction of each predictive set. For example, in the ciprofloxacin classifiers, a subset of genes belonged to the SOS regulon, consistent with the known induction of DNA damage repair pathways under fluoroquinolone treatment (Dörr *et al*, 2009). Similarly, components of the SoxR regulon, which regulates redox-responsive genes such as *mexGHI-opmD* (Sakhtah *et al*, 2016), were observed in classifiers for multiple antibiotics. These findings align with reports that antibiotic exposure triggers global stress response programs that facilitate bacterial survival (Dalbanjan *et al*, 2024; Dawan & Ahn, 2022; Niu *et al*, 2024).

However, the majority of selected genes did not map onto annotated operons or iModulons, suggesting that resistance-associated expression signatures extend beyond established regulons. This broad distribution highlights the multifactorial nature of resistance and the potential involvement of under characterized or strain-specific regulatory programs. Some markers may reflect cellular states shaped by prior antibiotic exposure, mutations in upstream regulators, or metabolic rewiring in response to treatment.

It is critical to distinguish between genes that contribute directly to resistance and those that serve as biomarkers of the resistant state. While several selected genes correspond to known resistance determinants, others may represent indirect indicators of resistance phenotypes without being mechanistically involved. For instance, upregulation of stress response genes may occur as a consequence of resistance-conferring mutations rather than serving as a primary defense. Accordingly, the interpretability of expression-based classifiers should be tempered by careful functional validation. Techniques such as gene knockouts or overexpression studies will be essential to confirm the roles of novel markers in antibiotic resistance.

In conclusion, our GA–AutoML framework has robustly identified minimal transcriptomic signatures that distinguish resistant from susceptible *P. aeruginosa* isolates with high accuracy. The predictive sets include both canonical resistance genes and novel, previously unannotated transcripts, revealing the complex and layered nature of bacterial adaptation to antibiotic pressure. Although a subset of these biomarkers map to known regulatory modules, most are broadly dispersed, suggesting that resistance involves a combination of direct mechanisms and global physiological adjustments. These findings offer new opportunities for diagnostic development and shed light on the systems-level dynamics underlying antimicrobial resistance.

## Methods

### Data Retrieval and Preprocessing

Gene expression and antibiotic resistance data for this study were obtained from the publicly available dataset published by Khaledi et al. (020) (Khaledi *et al*, 2020). The dataset comprises transcriptomic profiles and resistance phenotypes of 414 *P. aeruginosa* clinical isolates collected from diverse clinical and research institutions. The expression dataset was used to construct the feature matrix (*X*), while corresponding resistance classifications for the four antibiotics, formed the target vector (*y*). These datasets enabled the training of machine learning models to predict antibiotic resistance based on transcriptomic signatures.

Gene expression data were preprocessed to align with resistance labels, ensuring consistency in sample representation. Expression values were normalized, and genes with excessive missing values were excluded. Samples with incomplete resistance phenotype annotations were removed from the analysis.

### Threshold for Resistance Determination

Resistance and susceptibility classifications were assigned based on the minimal inhibitory concentrations determined in the original study (Khaledi *et al*, 2020). MIC values were binarized according to Clinical & Laboratory Standards Institute (CLSI) breakpoint criteria, with samples classified as resistant or susceptible for each antibiotic.

### Genetic Algorithm Optimization for Feature Selection

To systematically reduce the number of transcriptomic features while maintaining predictive accuracy, a genetic algorithm-based feature selection approach was implemented. GA iteratively refines feature subsets by mimicking the principles of natural selection, favoring gene sets that maximize predictive performance.

The GA pipeline was initialized with 1,000 randomly generated feature sets, each consisting of 40 genes. The selection of 40 genes was determined through an empirical evaluation process, where feature set sizes ranging from 16 to 40 genes were systematically tested using LR and SVM, with ROC-AUC and F1-score as primary evaluation metrics. Feature sets containing 40 genes consistently provided the best trade-off between predictive accuracy and feature set interpretability, making it the optimal configuration (Figure 2D-G). Each GA iteration refined these feature sets over 300 generations, systematically converging on the most informative subsets.

Two types of outputs were generated from the GA selection process: iteration-specific feature sets, representing the highest-performing gene subset identified in each iteration, and a ranked gene list, which quantified how frequently individual genes appeared in the top-performing subsets across all GA iterations. These outputs provided complementary insights, allowing both the selection of stable predictive markers and an assessment of the broader resistance-associated transcriptomic landscape.

To assess feature set stability, top-ranked gene lists from two independent GA runs were compared. The overlap between the highest-scoring gene sets in early runs was limited (∼30%), suggesting substantial variability in feature selection. To address this, the number of GA iterations were increased to 1000, and a ranked gene list was generated, quantifying the frequency with which specific genes appeared in the top-performing subsets across all iterations. The ranked gene list was further segmented into subsets of 20 to 50 genes and systematically evaluated to pinpoint the highest-performing gene sets.

A major limitation of GA-derived ranked lists was the presence of hypothetical genes, which reduced interpretability. To address this, iteration-specific feature sets were filtered to retain at least 80% of the well-characterized genes based on functional annotation databases. Although this filtering was also applied to the ranked gene list, evaluation shifted toward iteration-specific gene sets due to their superior classification performance. The final GA-selected gene subsets were validated externally and consistently maintained high precision and recall across all antibiotics (Table 1).

### Machine Learning Model Training and Evaluation

Automated machine learning was employed to optimize model training and hyperparameter selection. The AutoML pipeline was implemented using the auto-sklearn Python library, which systematically explores a range of models and hyperparameter configurations to identify the best-performing classifier for each antibiotic-specific dataset.

Preprocessed gene expression data were split into training (80%) and test (20%) sets while maintaining a stratified distribution of resistance phenotypes. To prevent overfitting, a stratified k-fold cross-validation approach (k=3) was applied during training. The AutoML classifier was configured with a runtime limit of five hours per model to ensure exhaustive search while maintaining computational feasibility. Feature sets derived from GA were provided as input, with model selection based on optimization of ROC-AUC and F1-score.

Model performance was assessed using multiple evaluation metrics to ensure robustness. ROC-AUC was used to measure the ability of the model to distinguish between resistant and susceptible strains, while F1-score provided a balance between precision and recall ensuring both classes were correctly classified. Precision quantified the proportion of correctly predicted resistant strains, whereas recall measured the fraction of actual resistant isolates correctly identified. Confusion matrix analysis was conducted to evaluate true positives, true negatives, false positives, and false negatives, providing further insights into misclassification patterns.

Final models were selected based on test set performance, and the best-performing classifiers were used for further downstream analysis.

### Independent Component Analysis and iModulon Mapping

To unravel the regulatory architecture underlying antibiotic resistance in *Pseudomonas aeruginosa*, we applied ICA using the iModulonMiner framework (Sastry *et al*, 2024). ICA decomposes transcriptomic data into independent components, termed iModulons, which represent co-regulated gene groups under shared regulatory control (Hyvärinen & Oja, 2000; Comon, 1994). This approach enables the identification of transcriptional regulatory networks driving resistance phenotypes.

We analyzed the normalized transcriptomic profiles of 414 *P. aeruginosa* PA14 clinical isolates, using the FastICA algorithm implemented in the iModulonMiner pipeline (https://github.com/SBRG/iModulonMiner). Input data consisted of a centered and scaled gene expression matrix (genes × samples). Stability analysis determined the optimal number of components to be 203. Each iModulon was characterized by two outputs: a gene weight matrix, quantifying gene contributions, and a sample activity matrix, representing iModulon activity across samples. Genes with absolute weights > 2.5 were considered significant contributors. Functional annotations of iModulons were assigned based on Gene Ontology (GO) terms and KEGG pathways, enabling insights into their biological roles.

The gene subsets identified via GA were mapped onto the derived iModulons to explore their regulatory basis. Distinct feature sets for different antibiotics were analyzed for their convergence on iModulons, reflecting potential regulatory mechanisms underlying resistance. Strain-specific iModulons unique to PA14 were also identified, highlighting novel regulatory adaptations. These analyses facilitated a deeper understanding of transcriptional regulation in the context of antibiotic resistance.

### Statistical Analysis and Visualization

All statistical analyses were performed in Python (v3.9). Machine learning and cross-validation metrics (e.g., F1-score, precision, recall) were computed using scikit-learn, while automated model selection was handled by auto-sklearn. All plots were created using matplotlib and seaborn, and final multi-panel figures were prepared using BioRender.

## Data Availability

Code and scripts for data analysis, machine learning, and visualization are available on GitHub at https://github.com/AAlsiyabi-Research-Group/Predicting-P.-aeruginosa-Resistance-with-Minimal-Gene-Signatures. Data sets are also provided.

## Author Contributions

N.S., S.A.S., and M.S. designed the study, preprocessed transcriptomic data, developed and implemented the GA–AutoML pipeline, conducted feature selection and classifier evaluation, performed biological interpretation, and wrote the manuscript. A.A.S. and R.S. supervised the study, provided conceptual guidance on methodology and biological validation, secured funding, and critically revised the manuscript. All authors reviewed and approved the final version of the manuscript.

## Acknowledgments

This research was supported by the National Institutes of Health (NIH) R35 MIRA grant (5R35GM143009) and the Nebraska Ethanol Board Award (26-1106-0157-001) awarded to Rajib Saha. Syed Ahsan Shahid acknowledges funding support from the University of Nizwa.

## Disclosure and Competing Interests Statement

The authors declare that they have no conflict of interest.

## Notes

### Competing Interest Statement

The authors have declared no competing interest.

